# Stable and robust differentiation programs are predominantly irreversible

**DOI:** 10.64898/2026.07.30.741799

**Authors:** Somya Mani, Gašper Tkačik

## Abstract

Multicellular differentiation programs are almost always irreversible, with cells rarely revisiting states observed during earlier developmental stages. An important factor contributing to such irreversibility may be the cellular context: the dependence of each cell’s fate decisions on inductive signals from other, simultaneously present cell types. Using a class of rule-based models for context-dependent cell fate decisions we examine the evolutionary basis for the observed prevalence of irreversibility. Our study suggests that irreversibility, instead of being directly selected for, could be widespread due to its association with two other key traits, stability and robustness, which are essential for reproducible, cancer-free organismal formation. We identify further structural properties of desirable fate-decision programs and illustrate how explicit knowledge of cellular context is required for predicting developmental outcomes.

---

During embryonic development, organisms undergo multiple transitions in cellular composition to ultimately arrive at their adult stage. At the cellular level, these organismal state transitions are defined by division and differentiation events at which individual cells multiply and adopt new, typically more specialized cell fates. When left undisturbed by injuries and other perturbations, cells rarely revisit fates encountered earlier during development. This property of cellular differentiation programs has previously been recognized as *irreversibility* [1]. While kinetic irreversibility of individual differentiation steps and its possible molecular basis have been studied previously [2], the broad prevalence of irreversibility at the level of entire developmental programs remains perplexing.

Irreversibility is well documented in animals, for example in the nervous system [3–5] and intestinal epithelium [6–8]. The degree to which plant development is irreversible is often questioned, since plant cell fate is governed predominantly by positional cues rather than by cell lineage, making the overall commitment to differentiated fates lower than in animals [9]. Irreversibility can nevertheless exist at the level of the entire differentiation program, without commitment at the cellular level: plant cells undergo highly characteristic sequences of signaling contexts during organ development, and irreversibility is easily apparent during the unperturbed development of roots [10], leaves [11], and stomata [12].

Cell differentiation trajectories are plastic on evolutionary timescales [13]. For instance, vertebrates evolved a novel developmental pathway towards the cartilage cell type [14], and developmental trajectories leading to xylem cells vary across angiosperm plants [15]. These trajectories can even be plastic within an organism’s lifetime: reversions of differentiation often occur during animal [16] as well as plant regeneration [17]. But such reversions are almost never a part of normal, unperturbed development. Different cell-intrinsic molecular mechanisms maintain irreversibility across both animals and plants, including chromatin modification [18, 19], coupling between cell cycle and differentiation [20, 21], and dedicated developmental gene regulatory network motifs that combine feed-forward and feedback interactions [22–25]. To our knowledge, there is only a single reported exception to the irreversibility of developmental programs, namely during fin development in teleost fish, where a reversion to a de-differentiated state is part of the normal progression [26].

Few theoretical studies have addressed the evolutionary basis of developmental program irreversibility. The best investigated case has been the germ-soma differentiation, relevant to the onset of multicellularity [27]. This case, however, does not explain the persistence of irreversibility – despite the evolutionary plasticity – of more complex differentiation programs. A computational survey of simulated cell differentiation programs suggested that tree-like differentiation maps are very unlikely *per se*; irreversibility should therefore be interpreted as a consequence of selection [28]. One suggested possibility is that selection for reproducible morphogenesis could drive the evolution of irreversible programs where mobile, dividing cells differentiate into stationary, non-dividing cells [29]. However, irreversible differentiation extends beyond morphogenetic roles, and development of complex tissues involves multi-stage sequences of precursor cell types whose differentiation is irreversible at each stage, for example during the development of mammalian pancreas [30] and during plant vascular development [31]. In another study, selection for large numbers of terminally differentiated cell types was shown to favor irreversibility [1]; however, context-dependence of cell fate decisions, which is necessary for the widespread prevalence of regeneration [16, 17], was not considered. Taken together, the existence of developmental plasticity on organismal as well as evolutionary timescales indicates that irreversibility of differentiation programs must be an evolvable trait. Yet we still lack a theoretical explanation for why this trait would be so widespread across the multicellular kingdoms of life, why dedicated molecular mechanisms would evolve to maintain it, and how it could co-exist with regeneration programs.

Here we take a rule-based modeling approach to address this knowledge gap. The central tenet we base our models on is the context-dependence of cell fate decisions: such decisions, informed by cell-cell communication mechanisms, will depend on the concurrent presence or absence of other cell types in a developing tissue. The rule-based modeling approach enables us to systematically navigate the landscape of developmental programs and computationally sample thousands of model instances. We screen for those that exhibit stability of cell numbers, i.e., do not show unbounded growth or collapse. Stability of cell number is a recognized property of many plant and animal systems [6, 32]; although plants continually initiate new organs throughout their lifetimes, cell numbers of a given type within each growth zone, such as the shoot apical meristem, are highly regulated [33]. We simulate normal developmental progression for stable programs as well as explore their robustness to mutations, perturbations, and cancers – insults that real organisms routinely face.

scriptsize

With these results in hand, we answer two important questions. First, we ask whether *cell differentiation maps* (CDMs) accurately summarize developmental programs [39]. As the most commonly used representation of cell fate choices, such maps are being reconstructed from single-cell transcriptomics data at scale and at an increasing pace [40]. We show that cell differentiation maps incur an essential information loss: in the presence of context-dependent fate decisions, such maps cannot even correctly predict qualitative developmental outcomes. Second, we explore the interplay between irreversibility, stability of cell number, and robustness. We show that irreversibility strongly correlates with stability and robustness, and analyze the sources of such correlation. Based on these results we hypothesize that the prevalence of irreversibility across multicellular life likely results from an indirect effect, via direct evolutionary selection for the stability and robustness of developmental programs.

## Context-dependent developmental programs

We model the developmental dynamics of organs or tissues in terms of their cell type composition, starting with a dedicated progenitor (examples in Table 1)^1^. For simplicity, we neglect any spatial structure of the tissue or the organ, focusing instead on the *developmental context* (Fig 1A): the binary “fingerprint” indicating whether any cells of a given type are currently present or absent (see SI Appendix Sec. 1). This grossly reduced representation assumes well-mixedness of the cellular population in which spatial relationships are not explicitly treated, an assumption most likely applicable for early developmental stages when the total number of cells is limited. Despite these drastic simplifications, we will soon see that the space of context-dependent developmental programs still scales super-exponentially with the number of possible cell types *N*, making exhaustive exploration difficult. On the flip side, our approach tracks all cell types regardless of whether they remain present at the adult stage of development or not, and will enumerate and simulate *all* cell fate decisions possible, whether they actually happen as part of the normal developmental process or are only triggered upon mutations and perturbations.

**Table 1.** Adult cell systems, their cell types, and embryonic progenitors.

| System | Adult cell types | Embryonic progenitors | Ref |
| --- | --- | --- | --- |
| lightgray Mammals: hematopoiesis | hematopoietic stem cells, all derived blood cells | hemogenic endothelial cells | [34] |
| Mammals: nervous system | adult neural stem cells, neurons, ependymal cells, astrocytes, oligodendrocytes | neuroepithelial cells | [35] |
| Fruit fly (*Drosophila*): gut | intestinal stem cells, enterocytes, enteroendocrine cells | endoblasts | [36] |
| Fruit fly (*Drosophila*): testis | hub cells, cyst stem cells (CySC), cyst cells | somatic gonadal precursors (SGP) | [32] |
| Flatworm (*Schmidtea*): body | pluripotent neoblasts, lineage primed progenitors, others | embryonic piwi-1+ cells | [37] |
| black Cress (*Arabidopsis*): body | shoot + root meristems, other adult differentiated cells | hypophysis, apical pole cell | [38] |

**Figure 1.**
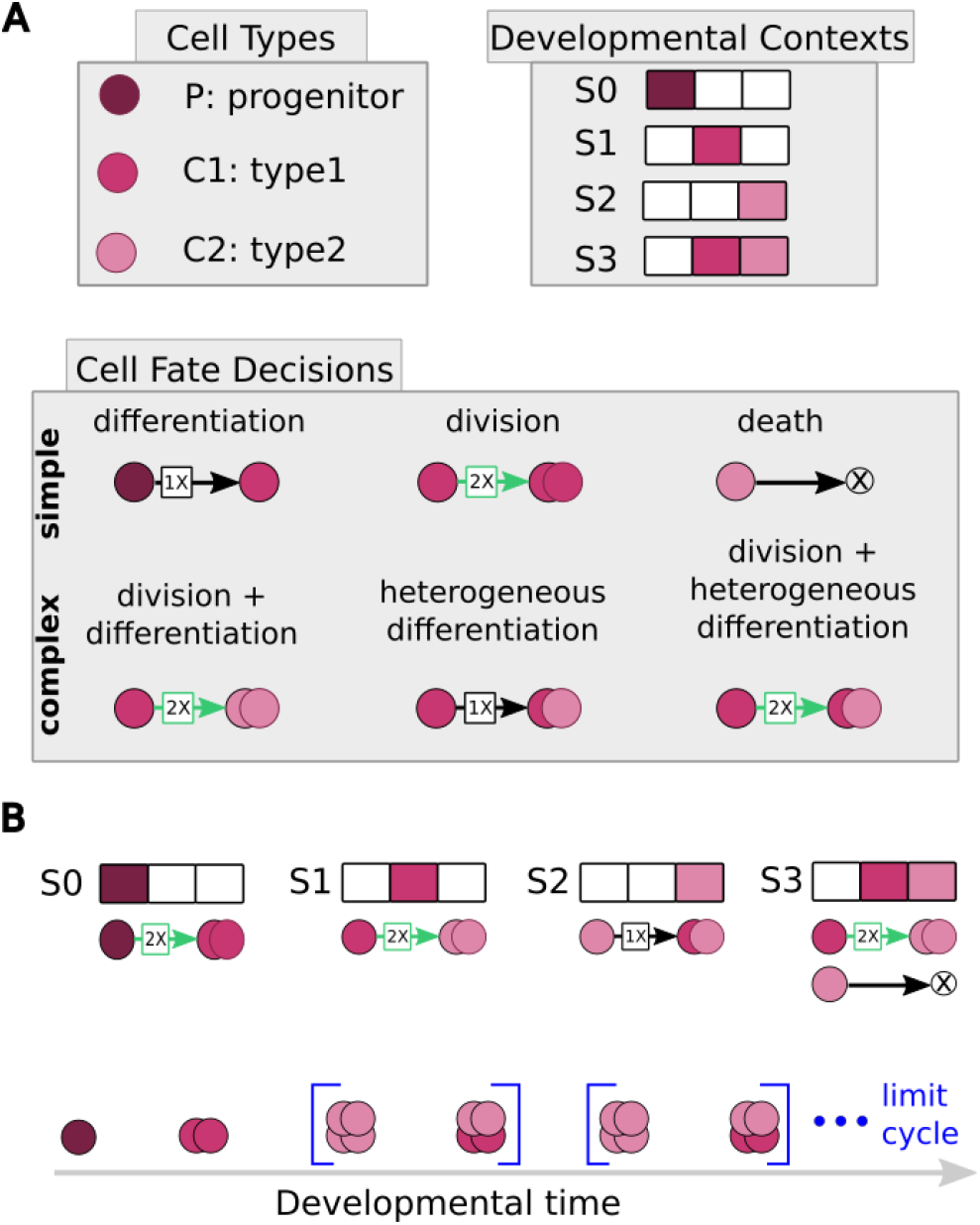
Model for context-dependent developmental programs. **(A)** Programs are specified by a list of cell types (in this illustrative example, *N* = 3 types: P = progenitor; C1, C2 = specialized types), giving rise to 2^3*−*1^ = 4 possible developmental contexts (S0-S3; presence/absence of cells of a given type is denoted by colored/white squares, respectively). Programs utilize six possible biologically-motivated cell fate decision rules (top row: *simple*; bottom row: *complex* ). Rules with black arrows do not increase the total cell count, but can decrease it (death) or change cell type counts (e.g., differentiation). Green 2X arrows denote divisions and thus increase total cell count; they can be coupled to simultaneous differentiation (see text). **(B)** *(Top*.) Example developmental program for a *N* = 3 system prescribes cell fate decision rule(s) that get triggered for every possible context. *(Bottom*.) Developmental dynamics obtained by applying the rules to the starting configuration with a single P cell present. After only two discrete time steps, this example system reaches a dynamical steady state (limit cycle), where the two configurations in blue brackets continually repeat.

We conceptualize developmental dynamics as a deterministic, discrete-time process, during which an organ or a tissue progresses from a single progenitor cell (= initial state) to its final composition (= stationary state), by invoking a fixed set of cell fate decision rules; we can think of the rules as genetically encoded, even though our framework does not treat such encoding explicitly. Each rule triggers during the timestep when its developmental context conditions are met (see SI Appendix Sec. 3). A stationary state represents the condition where the developmental context either reaches a fixed-point or a limit cycle even as cell numbers could keep changing. We track the dynamics of cell types (i.e., which types are present or absent at any given time) as well as cell counts (i.e., how many cells of each type are present).

Fate decisions are all formally treated as differentiation decisions, which may be preceded by a cell division in the same timestep. This approach easily accounts for simple cases: non-differentiation is represented by a decision to differentiate into the self cell type; cell death is represented by a decision to differentiate into a special, *null* type (⊗). The framework also accommodates complex cases motivated by biological observations: for instance, heterogeneous differentiation accounts for cases where cells of the same type partition equally between two fates [41]. In sum, our framework comprises six possible cell fate decisions, organized into three *simple* and three *complex* ones (Fig. 1A).

The set of all possible cell types and fate decision rules, along with their corresponding triggering developmental contexts, is called the *developmental program* (Fig. 1B) The space of possible developmental programs grows super-exponentially with the number of cell types, *N* (SI Appendix Fig. S1A). We use exhaustive enumeration (12 960 programs at *N* = 3) or Monte Carlo (26 400 programs at *N* = 4, 66 000 at *N* = 5 and *N* = 6) to sample and then directly simulate the programs and assess their stationary states in terms of the cell types and numbers present (Fig. 1B, see SI Appendix Sec. 2).

We call programs for which each of the *N* cell types appears at least once during development, even if not all are present in steady state, *non-trivial*. A tiny fraction of *invalid* programs that we exclude from consideration lead to pathological behaviors with fractional cell counts (SI Appendix Fig. S1B,C; Sec. 4)). The remaining *non-trivial, valid* programs can be categorized into three major classes based on their growth dynamics. For *shrinking* programs, the number of cells of all types keeps decreasing until the program terminates with zero cells. For *expanding* programs, at least one cell type keeps increasing in number indefinitely. For *stable* programs, non-zero long-term cell number is maintained once a steady state is reached; as we shall see, that steady state can either be a *fixed point* or a *limit cycle*. A small fraction (*<* 5%) of sampled cases give rise to more complex *transient cycle* dynamics, which we do not analyze further (SI Appendix Fig. S2). We also ignore *unicellular* outcomes that contain a single cell in their steady state (SI Appendix Fig. S3; Sec. 5). Taken together, our simulated data that we analyze below comprises 6 160 (*N* = 3), 6 144 (*N* = 4), 10 053 (*N* = 5), and 7 684 (*N* = 6) *non-trivial, valid, multicellular* programs, respectively.

To be concrete, we start with a small yet realistic developmental program depicted in Fig. 2A, which recapitulates *Drosophila* testis development, where cells specialize into hub cells (C1; magenta), as well as cyst stem (C2; pink) and cyst (C3; gray) cells, starting with the somatic gonadal precursor (P; violet; see Table 1).

**Figure 2.**
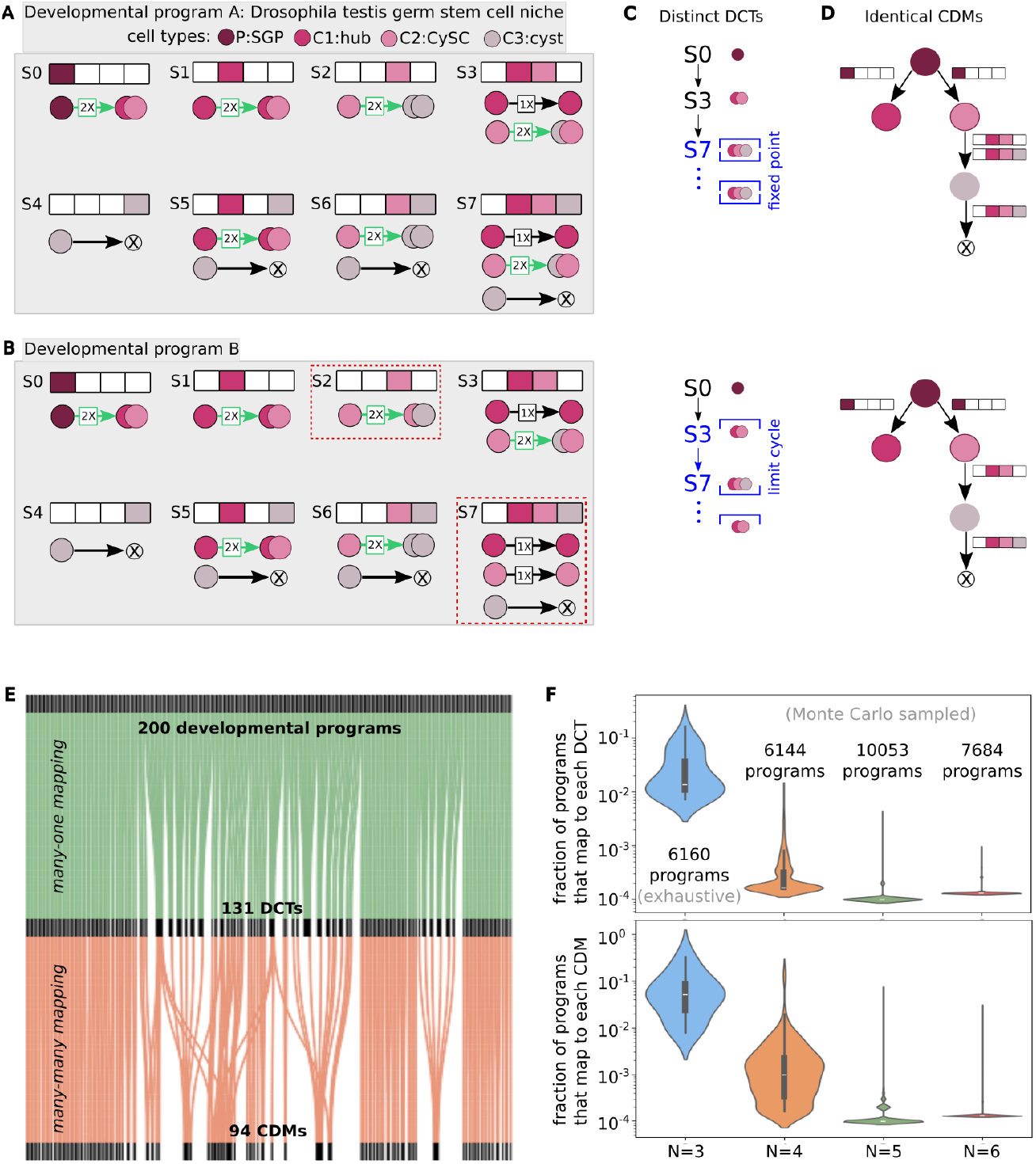
Cell Differentiation Maps are lossy representations of development. **(A,B)** Two example developmental programs for a *N* = 4 cell type organ, with cell types *P*, C1, C2, C3 and having 2^4*−*1^ = 8 possible contexts, S0 *−* S7. Program A corresponds to the *Drosophila* testis germ stem cell niche (see Table 1); Program B is a hand-crafted example which differs from Program A as highlighted by red dashed boxes. **(C)** Developmental Context Transition (DCT) backbones for Programs A (top) and B (bottom). Steady states are indicated with blue brackets: a [S7] fixed point for Program A, and a [S3, S7] limit cycle for Program B. **(D)** Programs A (top) and B (bottom) share identical Cell Differentiation Maps (CDMs). Note that a CDM is solely a graph of cell type relationships; here, we annotated the transition arrows by their corresponding developmental contexts to illustrate how identical CDMs can arise if context-dependence of transitions is ignored. **(E)** Mapping from developmental programs to DCT backbones (green lines) and further to CDMs (orange lines), depicted for a randomly chosen set of 200 *N* = 4 programs. Multiple programs can map to the same DCT, while the mapping between DCTs and CDMs is compressive and many-to-many. **(F)** Distributions (violin plots) of the fraction of sampled programs that map to a single DCT (top) or to a single CDM (bottom) for nontrivial, valid, multicelular programs with *N* cell types.

Cell fate decision rules have been extracted from published work [32]. We compare and contrast the behavior of this developmental program to a hand-crafted alternative (Fig. 2B), which contains the same cell types but slightly modifies two of the decision rules. Each program can be summarized by two graphs. A *Developmental Context Transition graph* (DCT) contains nodes that represent distinct contexts (Fig. 2C): a directed edge from node S to S^*′*^ implies that a cell fate decision rule which triggered in context S advanced the developmental state to context S^*′*^, whose cell type composition will be different if S^*′*^≠S. The sequence of such transitions starting at the initial state (progenitor cell P only) and ending at the stationary state is called the *DCT backbone* – this sequence represents contexts that are actually realized during normal development (see SI Appendix Sec. 6). Contexts (and their corresponding transitions) that are not part of the backbone are only realized when mutations and perturbations derail normal development.

A *Cell Differentiation Map* (CDM), in contrast, contains nodes that represent different cell types (Fig. 2D): a directed edge from node C to C^*′*^ implies that cell type C differentiates into cell type C^*′*^ (or into⊗ if cell death occurs from type C) at least once during normal development. Nodes in the CDM are ordered top-to-bottom (or left-to-right) according to the sequence in which cell types appear for the first time during development (see SI Appendix Sec. 7).

DCTs and CDMs both comprise compressed information about the original developmental program. CDMs are relevant because they are experimentally accessible [39, 40]. Backbone DCTs can be extracted from the original developmental programs, yet different programs could share the same backbone DCT; as we shall see, this could become relevant when we study deviations from normal developmental progression.

## Results

### CDMs are a lossy representation of development

We first wondered about the extent of information loss when complete developmental programs are replaced with their compressed representations. To assess this for CDMs, we ask how often unique CDM topologies are assigned to their corresponding programs. We find that the mapping from programs to CDMs is very lossy, which we demonstrate both by means of a hand-crafted example (Fig. 2A–D), as well as by an extensive statistical analysis (Fig. 2E,F). The reasons for this information loss are two-fold.

First, developmental programs specify cell fate decision rules for all possible developmental contexts – those that occur during normal development, as well as those that only manifest upon perturbations and mutations. While details vary, this statement would be true for real cells, independently of any modeling framework. DCTs and CDMs, however, only capture those transitions that actually occur (and were monitored) during normal developmental progression, at least given the current experimental as well as data-analysis paradigms. For example, developmental programs in Fig. 2A,B encode cell fate rules for all 8 contexts (S0*−*S7), while the corresponding DCTs and CDMs in Fig. 2C,D only describe transitions and differentiation for 3 of those 8 contexts, namely*{* S0, S3, S7*}*.

Second, CDMs are agnostic about the developmental context within which differentiation decisions occur. For example, CDMs for Programs A and B in Fig. 2D are identical even though they correspond to two distinct DCTs. In the hand-crafted example presented in the figure, the two identical CDMs summarize developmental programs whose long-term dynamical behaviors differ qualitatively: Program A settles to a fixed point while Program B converges to a limit cycle. Concretely, while both CDMs in Fig. 2D include the C2 *→* C3 differentiation, this decision occurs in program A during the S3 *→* S7 context transition as well as in the steady state, in S7. In Program B, in contrast, this fate decision occurs only during the S3*→* S7 transition. Taken together, this example makes abundantly clear that identical CDMs cannot discriminate even between qualitatively different developmental outcomes.

What is the scope of such information loss beyond a singular, hand-crafted example? We exhaustively analyze 6 160 *N* = 3 programs to find that they map to only 33 unique DCTs and 13 unique CDMs. Using Monte Carlo sampling, on average, 3.8, 1.4 and 1.1 distinct programs map to the same DCT, and 19, 1.9 and 1.2 distinct programs map to the same CDM, for *N* = 4, 5, 6 programs, respectively (Fig. 2E,F). Importantly, the distribution of CDMs across uniformly sampled developmental programs is extremely skewed: 2 out of 13 CDM topologies map to *>* 50% of *N* = 3 developmental programs, and 3 out of 322 CDM topologies map to *∼*40% of *N* = 4 developmental programs (SI Appendix Fig. S4A).

Taken together, our results highlight the interpretational limits of CDMs reconstructed from data: many developmental programs map to the same CDM even though their normal developmental progressions — and possibly also disrupted progressions in face of environmental insults – result in very different sequences of visited states, which can furthermore terminate with qualitatively different stationary outcomes. Because of the extremely skewed nature of the map from programs to CDMs, observing a given CDM can either imply a strong or a very weak constraint on the underlying developmental program, depending on the multiplicity of the map for a given CDM. Lastly and most importantly, these findings imply that any regularities observed in data-derived CDMs (for instance, the irreversibility of developmental programs that we focus on in this paper) must be statistically assessed for significance against a null expectation obtained by an unbiased sampling from the underlying space of developmental programs, rather than by any *ad hoc* sampling in the space of CDMs. This is the task we undertake below.

### CDMs of stable programs are irreversible

All birds, mammals, and most terrestrial arthropods – that together include many well studied model organisms – undergo *determinate growth*: a type of development where individuals reach a stable body size upon maturity [42]. Stable overall body size typically also implies a tightly controlled body plan in terms of cell types, their numerosity, as well as their organization across spatial scales.

Such stability imposes stringent requirements on developmental programs: cells must multiply as tissues and organs grow, yet such growth must ultimately become bounded and must terminate or become balanced by cell death to produce a healthy individual. In this vein, developmental context dependence of cell fate decisions provides a natural feedback mechanism whereby the current cellular composition informs individual cells to stop their excessive proliferation. This is illustrated in Fig. 3A for our Program A: by modulating the differentiation / division decisions in state S7, the developmental program can be switched from its original stable fixed point outcome, to shrinking (all cells die), expanding (run-away growth), or more complex dynamical outcomes. Importantly, however, the discrete nature of cells and their decisions ensures that stability need not be a fine tuned condition: as our model demonstrates, one bit (presence/absence) per cell type encoded in the developmental context is sufficient to inform which developmental rule to trigger so as to stabilize growth.

**Figure 3.**
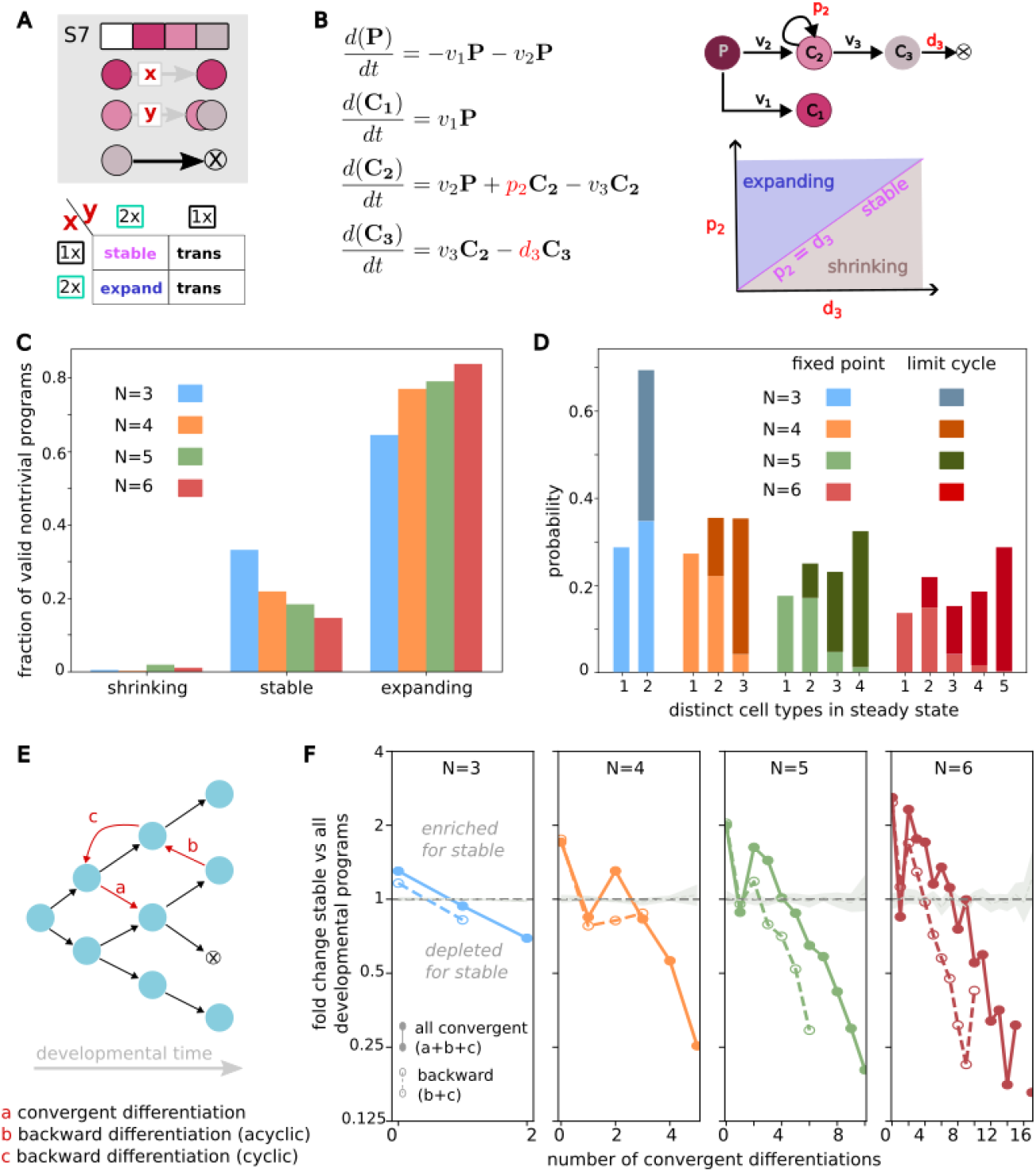
Stable programs are depleted for backward cell differentiations. **(A)** *(Top*.) Control over cell number stability for the Program A of Fig. 2, by varying cell fate decision rules for cell types C1 and C3 in steady state context S7. Stability of cell numbers depends on the nature of the gray arrows labeled x y. *(Bottom*.) A table summarizing long-term growth outcomes depending on whether x and y involve cell division: stability is achieved when the loss of C3 cells due to cell death is balanced by the division of C2. **(B)** *(Left*.) A standard, lineage-based ODE model for cell populations *c*1, …, *c*3 of the corresponding types C1, C2, C3, directly based on the Program A. *(Top right*.) Differentiation rates are given by parameters *v*0, …, *v*3 ; *p*2 is the division rate of C2; *d*3 is the death rate of C3. The ODE system has two steady states – one where C1, C2, C3 co-exist, and another with only C1. *(Bottom right*.) Coexistence of the three types with stable cell numbers, qualitatively matching the discrete dynamics of (A), only occurs on a finely tuned parameter manifold (violet line) that precisely balances division and death rates. **(C)** Histogram for the proportion of shrinking, stable, and expanding programs for different number of cell types, *N* . **(D)** Histograms for the number of distinct co-existing cell types at steady state of stable programs (light shade: fixed points; dark shade: limit cycle). **(E)** Schematic of a cell differentiation map (CDM) introducing different kinds of regular (black arrows) and convergent differentiations (red arrows). Blue circles: distinct cell types. Arrow (a) is a *forward convergent* differentiation; arrows (b) and (c) are *backward* differentiations. **(F)** Enrichment vs depletion of all convergent (solid) and backward convergent (dashed) differentiations in stable programs with *N* = 3, …, 6 cell types. Vertical axis shows the fold-change in the proportion of CDMs for stable vs all sampled developmental programs with a given number of convergent differentiations (horizontal arrow). Values *>* 1 indicate enrichment, values *<* 1 depletion. Gray shaded regions represent the sampling noise bands obtained through randomly shuffling the number of convergent differentiations for sampled programs and recalculating fold-change values. The shuffling was performed 1000 times and the shaded region represents standard deviations around mean fold-change values. Stable programs are enriched for zero convergent differentiations and strongly depleted for high number of convergent differentiations.

In contrast, standard lineage-based ordinary differential equation (ODE) models of development do not share this robust stabilization property. In Fig. 3B we construct the simplest such ODE model for Program A. Achieving a stable steady state where cell types C1, C2, C3 coexist, as they do in *Drosophila* testes and our discrete model, requires a finely-tuned balance between the rates of cell division, differentiation, and death that only occurs in a set of vanishing measure in the model’s parameter space. The stable co-existence region can be expanded by the addition of feedback loops [43], which – in contrast to our framework – operate with perfect information about the current cell composition^2^. While this mean-field approach should be applicable when the discreteness of individual cells and their decisions can be neglected at the tissue or organ mesoscale, it also intriguingly suggests that such discreteness may help, rather than hinder, achieving stable control over tissue and organ growth.

These considerations are reflected in a large-scale screen for stable developmental programs shown in Fig. 3C. We find that while only a small fraction (*<* 1%) of programs lead to shrinking dynamics and most lead to unbounded, expanding growth, a substantial fraction (*∼* 10 *−* 30%) are stable in cell type composition as well as in cell numbers. We observe a clear trend where stability is more difficult to achieve for programs that involve increasing numbers of cell types, likely because each type must be stable for the entire program to be stable. Real systems might mitigate this issue by combining developmental context dependence with global feedback, which is an interesting avenue for future research.

A majority of stable programs generate multiple cell types at steady state. These steady states can either be fixed points, consisting of a single developmental context, or limit cycles, where the program oscillates between several different contexts (Fig. 3D). Programs with a total of *N* cell types can generate a rich diversity of steady states that span the entire range: from organs that contain a single type in steady state (by definition, a fixed point), to organs where all *N* types appear in steady state (predominantly, as a limit cycle, as expected on general grounds [44]).

Cell Differentiation Maps (CDMs) are not directly informative about cell number stability. While they encode cell differentiation and death in their topology, they do not keep track of cell divisions. Consequently, the same CDM can correspond to expanding, stable, or shrinking programs. For example, 14% of the CDMs for *N* = 4 programs map to all three growth dynamics (SI Appendix Table. S1). Nevertheless, some CDMs are differentially enriched for programs with different growth outcomes, leading to statistical associations between CDM topology and stability (SI Appendix Fig. S4B).

The key association that we focus on here is that between *convergent differentiations* and stability. Convergent differentiations include all deviations from tree-like CDMs that can result in a given cell type being accessible from more than one parental type; among convergent differentiations, cyclic or acyclic *backward differentiations*, in particular, break the irreversibility of developmental programs (Fig. 3E). We find a strong and consistent depletion of convergent differentiations for stable developmental programs, those that produce fixed point steady states as well as those producing limit cycles, across the range of cell type numbers explored (Fig. 3F, SI Appendix Fig. S5). The abrupt depletion of CDMs with 1 convergent differentiation is driven by a handful of exceptionally popular CDMs which are predominantly produced by expanding developmental programs with limit cycle steady states (SI Appendix Fig. S6). The statistical association is particularly notable for backward differentiations, implying that selection for growth stability strongly favors irreversible, tree-like developmental programs.

### Robust developmental programs are irreversible and canalized

In nature, organisms develop from a zygote in the presence of many sources of noise and perturbations: environmental fluctuations and stresses, intrinsic stochasticity of cellular biochemistry, and impinging mutations all have the potential to derail the developmental process. An extensive body of work at the interface of evolution,systems, and developmental biology suggests that selection must have favored *robustness*, i.e., the property of developmental programs to reproducibly converge to a functional organismal outcome despite noise and perturbations [45, 46]. In the following, we analyze stable programs for three types of robustness – robustness to mutations that affect the developmental program, to perturbations such as injuries, and to cancer-causing somatic mutations – to evaluate if such robustness preferentially emerges in specific CDM topologies.

### Robustness to mutations

We model a mutation as a change in cell death or differentiation fate decision affecting a single cell type in a single context (Fig. 4A, red symbols). Mutations can divert developmental trajectories towards contexts not normally encountered during development, i.e., away from the DCT backbone, with a potential to disrupt the resulting steady state (Fig. 4B, red arrows; *m*1, *m*2 do not disrupt the steady state, while *m*3 does). To explore the consequences of mutations at scale, we randomly generate 100 independent mutants for each stable program (see SI Appendix Sec. 8.1). We define the *mutation robustness index* of a program as the fraction of mutants (out of 100 attempted) that map to the same steady state as the original, non-mutated program. Generally, stable programs are very mutationally robust: more than 70% (and for some *N* more than 90%) have a mutation robustness index *>* 0.7 (SI Appendix Fig. S7).

**Figure 4.**
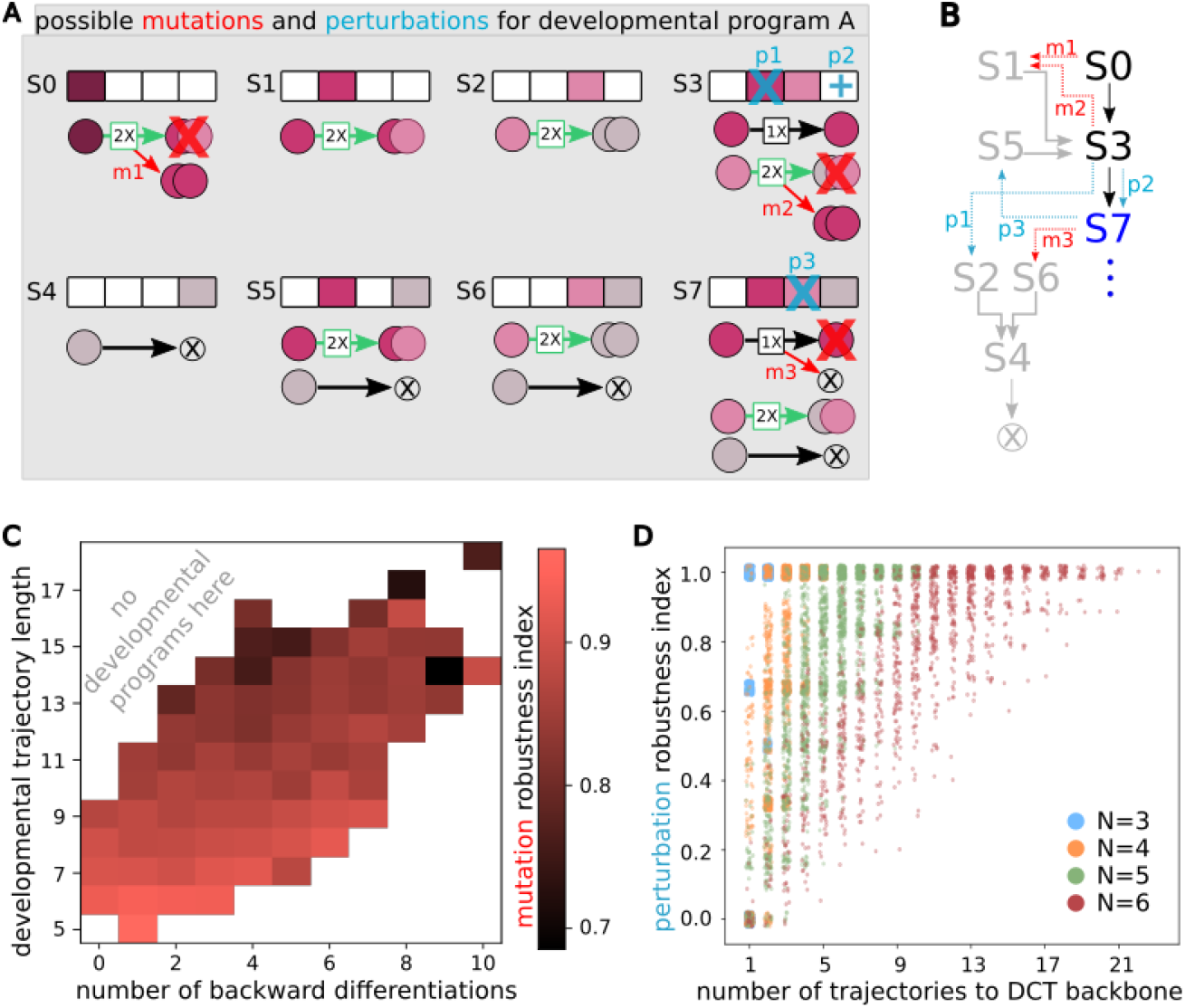
DCTs of robust programs are canalized and their CDMs are depleted for backward differentiations. **(A)** Examples of mutations (red symbols, *m*1, *m*2, *m*3) and perturbations (blue symbols, *p*1, *p*2, *p*3) for Program A from Fig. 2. Mutation *m*1 affects the differentiation decision of the progenitor; mutation *m*2 leads to erroneous cell differentiation of C2 in context S3; mutation *m*3 leads to erroneous cell death of C1. Perturbation *p*1 ablates all cells of type C1 from the developing organ when it is in context S3; perturbation *p*2 adds one cell of type C3 in context S3; perturbation *p*3 ablates all cells of type C2 in context S7. **(B)** Mutations and perturbations drive the developmental progression for Program A away from the DCT backbone (black arrows, trajectory length = 3), to access contexts encoded by the wild-type (non-mutant) program but never visited during normal development (gray arrows; trajectories starting in states S1, S2, S5, S6). Three trajectories, including the backbone, map to the non-perturbed steady state S7. Program A is therefore robust to mutations *m*1 and *m*2 (if they happen independently), but not *m*3, which would derail development via states S6 and S4 into cell death. Similarly, Program A is robust to perturbations *p*2 and *p*3 but not *p*1, which also leads to cell death via S2 and S4. **(C)** Average mutational robustness index (color; shown here for all 7 684 valid, nontrivial, multicellular *N* = 6 programs) is systematically higher for those programs that have shorter DCT backbone trajectories and smaller number of backward differentiations in their CDMs. **(D)** Perturbational robustness index (vertical axis) for all *N* = 3, … 6 programs (each dot = one valid, nontrivial, multicellular program) is higher for programs that are more *canalized*, i.e., have more DCT trajectories leading from non-backbone developmental contexts back to the DCT backbone.

We find that mutational robustness strongly and significantly correlates with short developmental trajectories. We quantify this by computing the Pearson correlation coefficient, *R*, between mutation robustness index and the length of the DCT backbone (with *R* = *−*0.43, *−* 0.70, *−*0.62, *−*0.54 for *N* = 3, 4, 5, 6 programs, respectively; all *p*-values *<* 10^*−*100^). Intuitively, because short developmental trajectories traverse fewer developmental contexts, they minimize the probability that a random mutation would impact a cell fate decision that is part of normal development. DCT backbone length also correlates with the number of backward convergent differentiations in the CDM (*R* = 0.39, 0.43, 0.63, 0.73 for *N* = 3, 4, 5, 6 programs, respectively; all *p*-values *<* 10^*−*60^). This in turn leads to negative correlations between mutational robustness and backward convergent differentiations which are otherwise only weakly related (partial *R* between mutation robustness index and backward convergent differentiations controlling for DCT backbone length are 0.02, 0.15, 0.07, 0.12 for *N* = 3, 4, 5, 6 programs, respectively; non-significant for *N* = 3, and *p*-values *<* 10^*−*60^ for *N* = 4, 5, 6). Thus, in our model, stable developmental programs that are mutationally robust tend to have short developmental trajectories and are predominantly irreversible (Fig. 4C, SI Appendix Fig. S9A).

### Robustness to perturbations

We model a perturbation as a spurious ablation or introduction of a cell type in a developing tissue or organ (Fig. 4A, blue symbols). Such perturbations can be caused by injuries or noise-induced errors in program execution. Like mutations, perturbations can also divert developmental trajectories away from the DCT backbone and can disrupt the resulting steady state (Fig. 4B, blue arrows; *p*2, *p*3 do not disrupt the steady state, while *p*1 does). To explore the consequences of perturbations at scale, we randomly generate 100 independent perturbations for each stable program, by ablating an existing cell type, or introducing a non-existing cell type, at a randomly chosen time step in the developmental progression (see SI Appendix Sec. 8.2). We define the *perturbation robustness index* of a program as the fraction of perturbations (out of 100 attempted) that map to the same steady state as the non-perturbed program. Generally, developmental programs are fairly perturbationally robust: more than 60% have a perturbation robustness index *>* 0.7 (SI Appendix Fig. S8).

We find that perturbation robustness strongly and significantly correlates with the number of trajectories that start off the DCT backbone yet lead back towards it, thereby ensuring a return to the original steady state upon perturbation (*R* = 0.48, 0.70, 0.75, 0.80 for *N* = 3, 4, 5, 6 programs, respectively; all *p*-values *<* 10^*−*100^). This bias is independent of irreversibility, and grows increasingly prominent with *N* (Fig. 4D, SI Appendix Fig. S9B), likely because the number of developmental contexts that are not on the DCT backbone yet can be induced by a perturbation grows exponentially with *N* . Thus, in our model, stable programs that are perturbationally robust tend to become *canalized* [47]: they feature a DCT backbone which is an attractor of developmental dynamics [48].

### Robustness to cancer-inducing somatic mutations

A somatic mutation is incident upon a single cell in the developing tissue or organ and leaves the rest of the cells unaffected. Somatic mutations, especially those that disrupt cellular signaling, can result in cancers [49]. This typically occurs because a mutated cell continues to proliferate due to a failure to correctly sense and interpret signals that would otherwise instruct it to stop. Here, we recapitulate this phenomenology by simulating somatic mutations which affect a cells’ ability to correctly parse their developmental context. A biologically plausible scenario could assume, for example, that each cell displays signals which indicate its own type, and carries receptors which allow it to parse the identity signals from other cells; these signals are decoded into the cell’s fate decision rules by a possibly complex *receptor logic* that we do not model explicitly. Somatic mutations we consider disrupt this receptor logic and lead to a mismatch between the actual and perceived contexts (Fig. 5A). Concretely, we simulate the effect of a somatic mutation by replacing, in steady state, a single, randomly chosen, mutated cell’s actual contexts that govern its cell fate decision rules with uniformly randomly chosen perceived contexts, keeping intact only the ability of the cell to self-identify (Fig. 5B).

**Figure 5.**
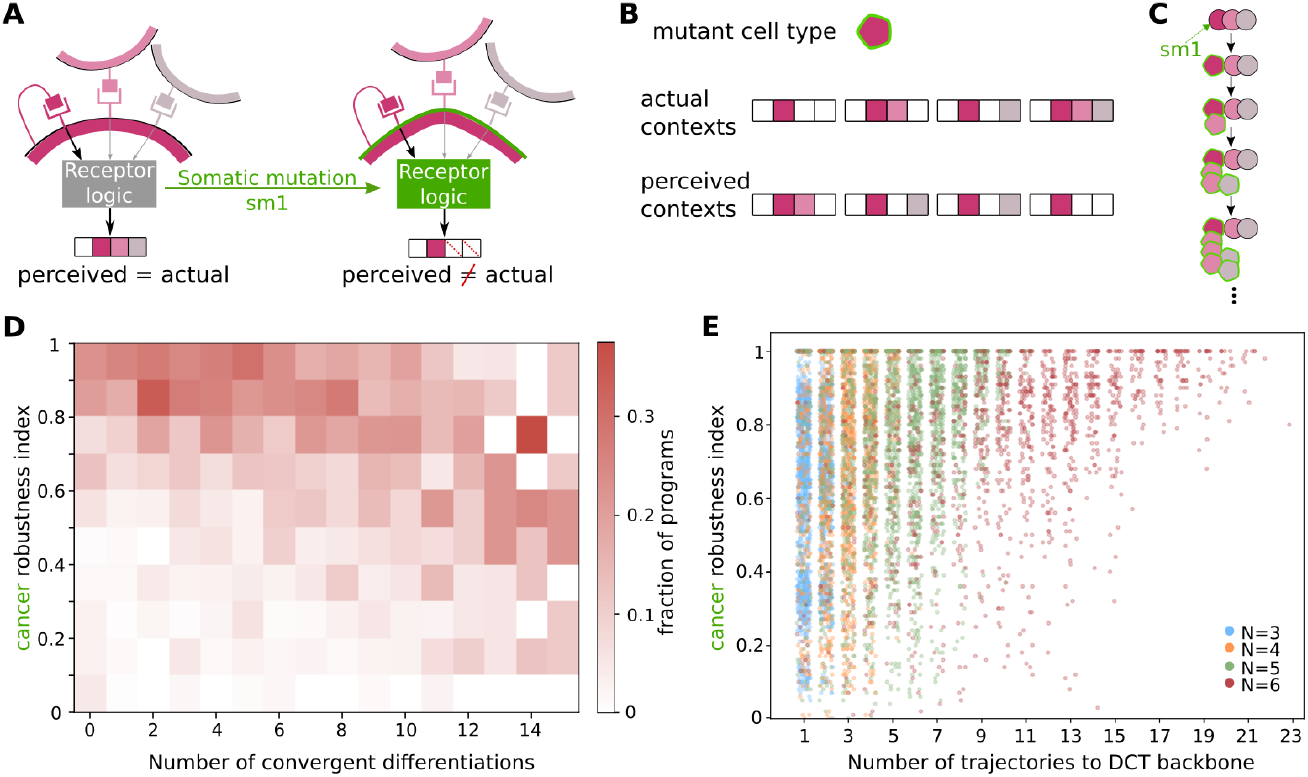
Developmental programs robust to cancer are canalized and their CDMs are depleted for backward differentiations. **(A)** Somatic mutations cause cells to incorrectly perceive their developmental context. *(Left*.) A normal cell of type C1 receives signals from itself (magenta) and from cells of types C2 (pink) and C3 (gray). *(Right*.) A somatic mutation alters the *receptor logic* such that the cell perceives the context incorrectly and thus possibly triggers inappropriate decision rules. In this example, somatic mutation *sm*1 causes a cell of type C1 in context S7 to incorrectly infer that the context is S1. **(B)** Possible effect of a somatic mutation on cells of type C1. After a mutation, the actual context of a mutated cell is erroneously perceived as shown, by replacing its actual contexts by random draws from all possible contexts (while keeping the ability of cells to self-identify intact). **(C)** Somatic mutation *sm*1 causes cancer when it is incident on a single cell of type C1 in the steady state S7 of Program A of Fig. 2A, because it leads to an uncontrolled growth of the mutated C2 population. **(D)** Cancer robustness index (vertical axis) is systematically higher for those programs that have fewer convergent differentiations in the CDM (horizontal axis). Shown is a conditional probability distribution (color) for cancer robustness index given the number of convergent differentiations (e.g., columns sum to 1), across all 7 684 valid, nontrivial, multicellular *N* = 6 programs (color). **(E)** Cancer robustness index (vertical axis) for all *N* = 3, … 6 programs (each dot = one valid, nontrivial, multicellular program) is higher for programs that are more *canalized*, i.e., have more DCT trajectories leading from non-backbone developmental contexts back to the DCT backbone.

Somatic mutations are inherited by all daughter cells of the initial mutant cell even as they differentiate and proliferate, potentially giving rise to a catastrophic cancer-like failure in which an original stable steady state of developmental dynamics is transformed into unbounded growth (Fig. 5C). This property makes the exploration of somatic mutations qualitatively different from the (germline) mutations in developmental programs and perturbations, which we described previously. To explore the consequences of somatic mutations at scale, we randomly generate 100 independent somatic mutations for each stable program incident on individual, randomly chosen cells in the steady state (see SI Appendix Sec. 9). We define *cancer robustness index* of a program as the fraction of somatic mutations (out of 100 attempted) that do not lead to unbounded growth. Generally, developmental programs are fairly robust to cancer: *∼* 80% of *N* = 3, …, 6 have a cancer robustness index*>* 0.5 (SI Appendix Fig. S10).

We find that cancer robustness index modestly but significantly negatively correlates with the number of convergent differentiations in the CDM (*R* = *−*0.01; *p*-value = 0.72 for *N* = 3 and *R* =*−* 0.21, *−* 0.34, *−* 0.31 for *N* = 4, 5, 6 programs, respectively; *p*-values *<* 10^*−*20^), as depicted specifically for *N* = 6 in Fig. 5D). Interestingly, cancer robustness correlates with the number of trajectories that start off the DCT backbone yet lead back towards it, analogously to the case of perturbational robustness (*R* = 0.06; *p*-value=0.005 for *N* = 3 and *R* = 0.25, 0.37, 0.48 for *N* = 4, 5, 6 programs, respectively; *p*-values *<* 10^*−*30^). This trend again grows increasingly prominent with *N* (Fig. 5E). Thus, in our model, stable programs robust to cancers tend to be canalized and irreversible.

## Discussion

In this study, we present a rule-based model of development where cell fate decisions depend on cellular context. We use this framework to generate thousands of artificial developmental programs in order to understand the evolutionary basis for certain commonalities in developmental processes prevalently observed in nature. In particular, we ask why multicellular differentiation programs tend to be irreversible, with cell differentiation maps that are tree-like. Our results indicate that irreversibility is strongly associated with stability of cell numbers and robustness to mutations, perturbations and cancers – traits that are functionally important, and thus possibly under selection, in multicellular organisms. Our work provides a rationale for why irreversible differentiation programs are so widespread across multicellular organisms despite multiple evolutionary origins of multicellularity, while cell-type switching is predominantly reversible for unicellular organisms, even for complex protists closely related to animals [50].

During development, cells make fate decisions to divide, die, or differentiate, depending on inductive cues from their neighbors. Our analysis demonstrates the extent to which cell differentiation maps, being agnostic about such cues and lacking the information about developmental context, are insufficient for predicting developmental progression and outcomes. It is important to be aware of such limitations given that cell differentiation maps are an increasingly common output of single cell transcriptomic studies [40]. Going forward, we envision that these limitations could be remedied by experimental advancements in spatial transcriptomics which measure both cellular transcriptomes and cellular contexts [51], and by methods such as the CellPhoneDB toolkit [52], which infer cellular interactions within developing tissues.

In addition to being agnostic about developmental context, cell differentiation maps carry no information about cell divisions. Perhaps surprisingly therefore, we show that their topologies can nevertheless be statistically linked to growth dynamics. Specifically, we find that stable developmental programs are depleted for backwards differentiations, suggesting that irreversible differentiation programs of real organisms could be due to their determinate growth dynamics. Conversely, our simulations show that expanding programs tend to have reversible differentiation programs. Although it might be tempting to extend this simulational finding to natural organisms with indeterminate growth, doing so could be unwarranted: The model’s expanding programs are a mixed-bag and include a variety of growth patterns, such as constant linear growth or exponential growth. On the other hand, cell divisions and deaths in real organisms are highly regulated, for example, in the gills of teleost fish [53], and in the shoot apical meristems of plants [33], making it inappropriate to compare real organisms with the whole class of model-generated expanding programs. Additionally, It is unclear if our framework can properly capture such regulation, which could occur via global growth feedback loops and not only through the developmental context, such as happens in the case of the sea anemone *Nematostella vectensis* [54].

Unlike growth stability, which is a property of individuals, biological robustness is a variational property [55]: over evolutionary timescales, developmental programs have accrued constraints that safeguard organisms against dysfunctional or disease states. These constraints are often *cryptic*, i.e., they involve organismal states that are rarely observed during the normal course of development, but manifest themselves, and may be essential in the response to, perturbations and mutations. Such cryptic states are typically not reflected in cellular differentiation maps, or in the backbone of developmental state transitions, as both of these abstractions are lossy descriptions of complete developmental programs. Since our approach involves sampling developmental programs directly, we were able to study cryptic constraints and variational properties such as different types of robustness. We focused on three types of universal insults that multicellular organisms face, independently of their evolutionary lineage: germline mutations, perturbations, and cancer-inducing somatic mutations. We find that developmental programs robust to these three types of insults are canalized and predominantly have irreversible differentiation programs.

Taken together, stability and robustness provide two independent, universal, and potentially selectable traits that go hand-in-hand with the irreversibility of developmental programs. Irreversibility, *per se*, seems nontrivial and should not be viewed as a default expectation [1, 28], although it is difficult to make this statement rigorous. This is because, on the one hand, mutational bias estimates for such a systems-level property are currently empirically out of reach, as is the resulting quantitative neutral expectation for “irreversibility”. On the other hand, we also highlighted statistical difficulties arising due to the skewed relationship between cell differentiation maps and the underlying developmental programs. Given these considerations, the only approach at our disposal was a computational study that clearly comes with its own assumptions and idiosyncrasies, which we fully acknowledge. Within these assumptions, however, irreversibility is uncommon and – unless it were directly selected for, for reasons that are not apparent from the current literature – could arise via indirect selection, with evolution favoring either stability or robustness, or both traits jointly.

In addition to the major methodological assumption highlighted above, we discuss two further limitations of our study. First, our simulations rely on drawing independent and uniform samples of developmental programs, allowing us to explore what Jacob referred to as the space of “the Possible” [56]. This independence fundamentally differs from “the Actual,” namely the developmental programs of organisms observed in nature, which are related by descent. Therefore, similarity of cell differentiation maps between phylogenetically close organisms is likely a consequence of similar underlying developmental programs; evolutionary relatedness could thus mitigate the effects of information loss in CDMs, which our model cannot capture.

Second, while context dependence of cell fate decisions is a universal property of multicellular development, our approach ignores the cellular and molecular mechanisms that implement this property, e.g., developmental signaling and gene regulatory networks (GRNs). This choice comes with trade-offs: many GRNs can produce the same developmental program, so our abstraction allows us to efficiently sample the huge space of possible programs. Furthermore, because our results are statistical and not mechanistic in nature, multiple mechanisms might lead to the same overall association between irreversibility and stability or robustness across different developmental programs. On the other hand, our choice makes it difficult to connect to data. Direct experimental elucidation of developmental programs is challenging and rare, but there is considerably more information assembled about the structure of real developmental GRNs, which we currently cannot utilize. Our model also does not capture how organisms with different GRNs, even if they encode the same developmental program, differ in the space of heritable variation that is available to them. In future work, adding a GRN layer to the model would allow us to better connect to well-characterized GRNs, e.g., for *Caenorhabditis elegans* [57] or *Arabidopsis thaliana* [58]. Taken together, grounding context-dependent fate decisions in an explicit GRN architecture could help us bridge the gap between the statistical patterns we identify here and the underlying molecular mechanisms that are the focus of many contemporary studies.

## Acknowledgments

SM was supported by a Resident Fellowship at the Konrad Lorenz Institute for Evolution and Cognition Research. We thank Yuriy Pichugin, Barbara Fischer, Pascal Hagolani and Günter Wagner for helpful discussions.

## Footnotes

1 Taken to the extreme, our modeling framework could apply to the entire organism with the embryonic progenitor being the zygote itself.

2 An analogous exploration to ours in the full space of lineage-based ODE models with feedback would have been intractable, as there is no convenient and systematic way to enumerate all developmental programs.

