## Supplemental Text & Figures for "Stable and robust differentiation programs are predominantly irreversible"

#### 1 State of an organ and developmental context

We model the development of organs with $N = \{3, 4, 5, 6\}$ cell types. These cell types include cell types in the adult as well as embryonic cell types. The state of an organ in the model is given by an $N$-length vector $\Omega$, where $\Omega_t^i$ is the number of cells of cell type $i \in \{1, 2, ..N\}$ in the developing organ at time-step $t$. Cell type 1 (`C1`) always represents the progenitor, and it never co-occurs with any other cell type.

The developmental context of the organ is given by an $N$-length binary vector $\kappa$:

$$\begin{aligned} \kappa_t^i &= 1 \quad \text{if } \Omega_t^i > 0 \\ &= 0 \quad \text{if } \Omega_t^i = 0. \end{aligned}$$

$\chi_t$ is the standard binary representation of the developmental context $\kappa_t$ of the organ at time-step $t$. Considering the constraint on the progenitor cell mentioned above, an organ with $N$ cell types can be in one of $2^{N-1}$ possible developmental contexts, and $\chi_t \leq 2^{N-1}$.

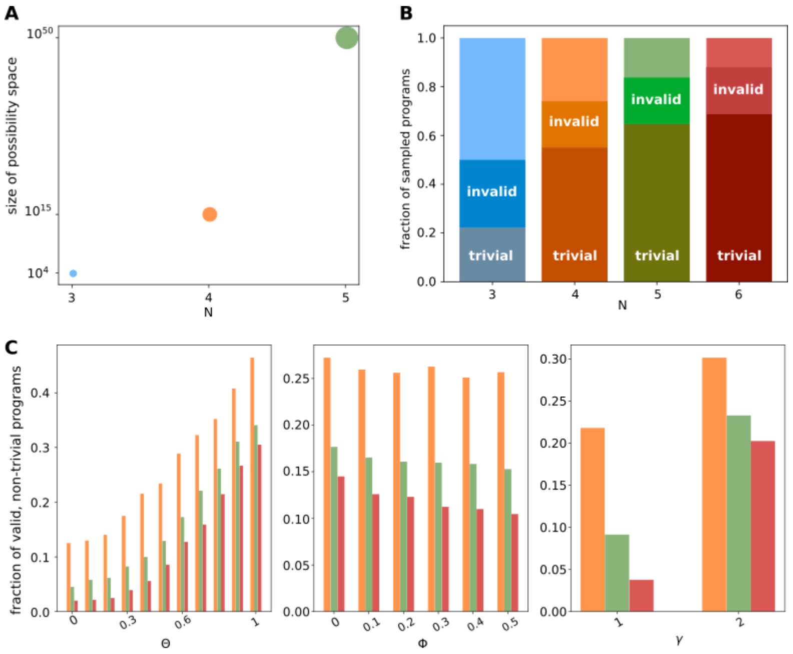


Fig. S1: Non-trivial and valid developmental programs. (A) Number of all possible $N = 3, 4, 5$ developmental programs. Dot sizes indicate relative sizes of the corresponding spaces. (B) Histograms showing the fraction of sampled programs that are trivial, non-trivial but invalid, and non-trivial and valid for $N = 3, 4, 5, 6$. (C) Histograms showing the fraction of programs sampled using a given value of a parameter (x-axis) that were non-trivial and valid: (left) cell division parameter $\theta$, (center) cell death parameter $\phi$, (right) parameter controlling heterogeneous cell differentiation, $\gamma$.

#### 2 Sampling developmental programs using ensemble parameters

We use a Latin hypercube sampling method to sample ensembles of deterministic *developmental programs* [1]. The ensembles are characterized by 3 parameters: $\theta$ is the average number of cell types whose cells divide in a given *developmental context*, $\phi$ is the average number of cell types whose cells die in a given *developmental context*, and $\gamma$ is the maximum number of distinct target cell types that cells of some mother cell type can differentiate into in a given *developmental context*. Briefly, we use these three parameters to split the space of all possible developmental programs into a uniform 3D grid, and sample uniformly within each grid square. In the following, we describe how we randomly generate programs using parameter values corresponding to each grid square:

Cells in the model can divide, die and differentiate. Each cell in an organ reads the developmental context in order to make these cell fate decisions. Two constraints apply: First, `C1` represents the progenitor, and no cell type, including itself, is allowed to differentiate into it. Second, developmental programs which include developmental contexts under which all cell types composing an organ to die are not allowed. Developmental programs in the model are encoded in terms of two sets of rules: $Div$ and $Diff$.

$Div$ is a set of $2^{N-1}$ N-length vectors

$$\begin{aligned} Div^i_{\chi_t} &= 2, \quad \Rightarrow \text{cells of cell type } i \text{ divide in context } \chi_t \\ &= 1, \quad \text{otherwise} \end{aligned}$$

The parameter $\theta$ is used to generate cell division rules:

$$\begin{aligned} Div^i_{\chi_t} &= 2 \text{ with probability } \theta \\ &= 1 \text{ with probability } 1-\theta \end{aligned}$$

$Diff$ is a set of $2^{N-1}$ matrices of size $N \times N$. $Diff_{\chi_t}$ gives the cell differentiation rules for an organ in a context $\chi_t$. $Diff$ is subject to the constraint that no cell type is allowed to differentiate into cell type-1, which is the progenitor.

$$\begin{aligned} Diff^{i,j}_{\chi_t} &> 0 \text{ if cells of cell type } i \\ &\qquad \text{differentiate into cell type } j \\ &= 0, \quad \text{otherwise} \end{aligned}$$

Additionally, if a cell type $i$ is programmed to undergo cell death in context $\chi_t$, then $Diff^{i,k}_{\chi_t} = 0 \ \forall k \in \{1, 2, .., N\}$.

Parameters $\phi$ and $\gamma$ are together used to generate $Diff$: Let $targets^i_{\chi_t}$ be the number of distinct cell types any cell of cell type $i$ can differentiate into in the context $\chi_t$. $targets^i_{\chi_t} = 0$ with probability $\phi$. Otherwise, $targets^i_{\chi_t} = 1$ if $\gamma = 1$, and $targets^i_{\chi_t} = 1$, or 2 with equal probability for $\gamma = 2$.

We choose the identities of the $targets^i_{\chi_t}$ cell types randomly from the set $\{1, 2, .., N\}$ with uniform probability. Let the set of target cell types for a cell type $i$ be $J$. Then,

$$\begin{aligned} Diff^{i,j}_{\chi_t} &= \frac{1}{targets^i_{\chi_t}} \forall j \in J \\ &= 0, \ \text{otherwise} \end{aligned}$$

We exhaustively sample all 12960 programs for N=3. For N=4, we sampled 200 independently generated random programs for each combination of ensemble parameter values; i.e. $11 \times 6 \times 2 \times 200 = 26400$ programs. Since the space of possible programs expands super-exponentially with N (Fig. S1A), in order to better sample the much larger space of programs for N=5,6, we generate 500 independent random programs for each combination of model parameter values; i.e., we sample $11 \times 6 \times 2 \times 500 = 66000$ programs. Overall, our dataset consists of $12960 + 26400 + 66000 + 66000 = 171360$ developmental programs.

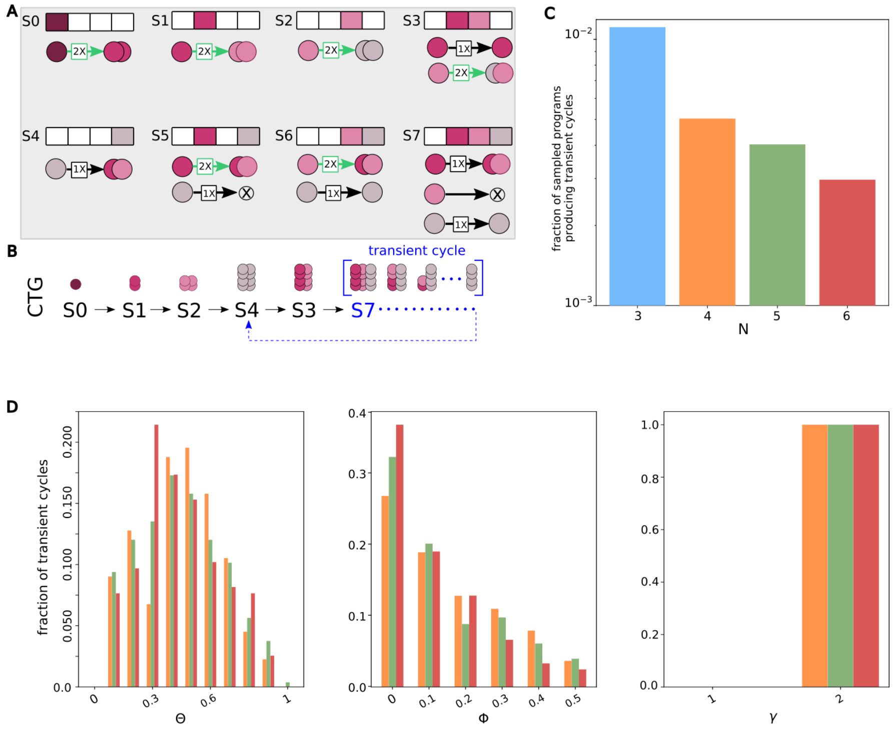


Fig. S2: Steady states with transient cycles. (A) Example developmental program that produces a steady state with a transient cycle at S7. (B) developmental dynamics for program in (A). (C) Bar-plot showing the fraction of sampled programs producing transient dynamics. (D) Fraction of programs with transient dynamics sampled using parameter values indicated in the x-axis for (left) cell division parameter $\theta$, (center) cell death parameter $\phi$, (right) parameter controlling heterogeneous cell differentiation, $\gamma$.

### 3 Updating the state of an organ

In the initial state, organs contain a single progenitor cell, i.e. $\Omega_0^1 = 1$ and $\Omega_0^i = 0,\ \forall i > 1$. We update the state of an organ in discrete time-steps. In each time-step, each cell in the organ makes cell division, cell death and cell differentiation decisions, in that order.

Let $\Theta_t$ be the state of the organ at time-step $t$ directly after cell-division. $\Theta_t^i = \Omega_t^i \times Div_{\chi_t}^i$.

And,

$$\Omega_{t+1}^i = \sum_k \Theta_t^k \times Diff_{\chi_t}^{k,i}$$

We update the state of the organ until a time-step $t_{ss}$ at which a previously encountered developmental context repeats; i.e., $\chi_{t_{ss}} = \chi_{t_{ss}-p}$. Since cells make cell fate decisions based on their developmental contexts, cells at time-step $t_{ss}$ make the same cell fate decisions as do cells at $t_{ss} - p$, and the sequence of developmental contexts$\chi_{t_{ss}-p}, \chi_{t_{ss}-p+1}, ..., \chi_{t_{ss}}$ repeats in a cycle. This cycle represents the steady state of the organ and $p$ is the period of the oscillation. organs that arrive at a steady state which is a fixed-point have $p = 1$. Since the number of possible distinct developmental contexts is $2^{N-1}$, all organs arrive at their steady state at $t_{ss} \leq 2^{N-1}$.

Note that while the developmental contexts $\chi$ repeat in a cycle, the states of the organ $\Omega_{t_{ss}-p}$ and $\Omega_{t_{ss}}$ are not necessarily the same, since number of cells of different cell types can still be changing across the cycle. Based on the dynamics of cell numbers, we classify programs into the following categories:

- Stable: $\Omega_{t_{ss}}^i = \Omega_{t_{ss}-p}^i\ \forall i \in \{1, 2, ..., N\}$.
- Expanding: $\Omega_{t_{ss}}^i \geq \Omega_{t_{ss}-p}^i\ \forall i \in \{1, 2, ..., N\}$
- Shrinking: $\Omega_{t_{ss}}^i \leq \Omega_{t_{ss}-p}^i\ \forall i \in \{1, 2, ..., N\}$
- Transient cycles: In a small minority of programs ($< 1\%$), in the steady state, the number of some, but not all, cell types shrink. In such cases, the shrinking cell types are eventually depleted after a few rounds of the steady state cycle and the organ jumps to a new state devoid of these shrinking cell types (Fig. S2). Since these do not capture any natural phenomena, we do not analyze such programs in this study.

### 4 Validity and non-triviality of developmental programs

- Validity: We call a set of developmental programs *valid* when $\Omega_t^i \geq 1$ for all cell types $i$ that are present at time-step $t \leq t_{ss}$. This is to disallow unrealistic rules that involve organs that contain a fraction of a cell of some cell type (Fig. S1B,C).
- Non-triviality: Let $\alpha$ be an $N$-length vector such that $\alpha^i = \sum_{t=0}^{t_{ss}} \chi_t^i$. That is, $\alpha^i > 0$ for all cell types that have appeared during the development of the organ. We call a set of developmental programs *trivial* when fewer than $N$ cell types appear during development of an organ, since this organ could have been produced using developmental programs for a smaller number of cell types (Fig. S1B,C).

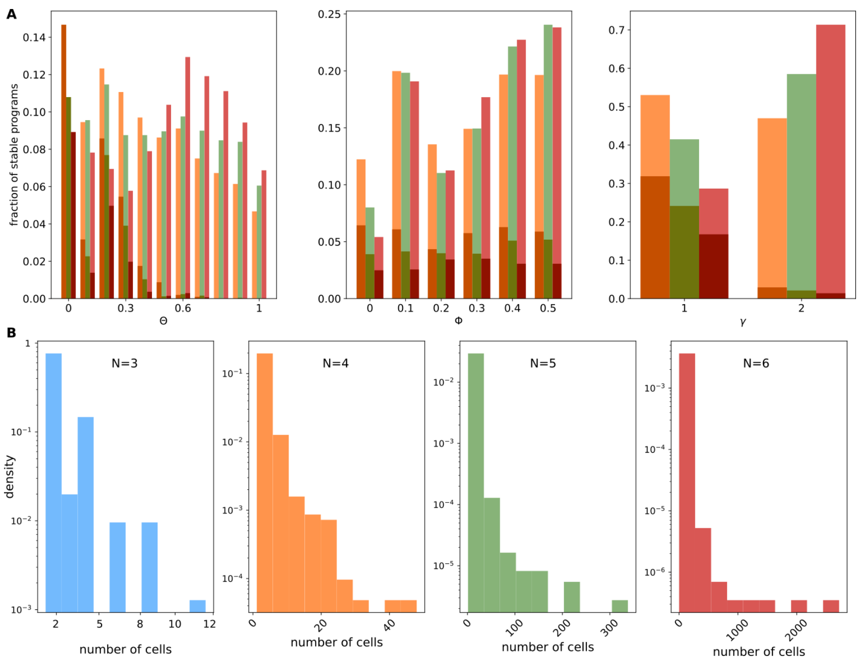


Fig. S3: Multicellular stable programs and the number of cells in their steady states. (A) Histograms showing the fraction of stable programs sampled using a given value of a parameter (x-axis) that were unicellular (darker colors), or multicellular (lighter colors): (left) cell division parameter $\theta$, (center) cell death parameter $\phi$, (right) parameter controlling heterogeneous cell differentiation, $\gamma$. (B) Histograms for the number of cells in N=3,4,5,6 multicellular stable programs sampled in this work.

### 5 Properties of stable programs

We further characterize stable programs as *unicellular* programs; these have a single cell in the final steady state, and *multicellular* programs; these have $> 1$ cell in the final steady state, even if the steady state consists of a single cell type. Unicellular programs are more likely to be generated at low values of the cell division parameter $\theta$ and when the heterogeneous differentiation parameter $\gamma = 1$ (Fig. S3A).

Multicellular stable programs vary widely in the number of cells in the adult stage: this is the total number of cells in the fixed point steady state, or the sum of all cells across all phases of a limit cycle steady state. The maximum number of possible cells in a stable organism is intrinsically related to the number of cell types, N – we simulate development using deterministic dynamics, therefore the maximum length of a developmental trajectory is bounded by the number of possible contexts (i.e., $2^{N-1}$). The largest possible organs are those whose cells divide in each context, thus the largest organs have $2^{2^{N-1}}$ cells. Most multicellular stable programs tend to be much smaller than this limit (Fig. S3B).

### 6 Developmental Context Transition Graph

The Developmental Context Transition Graph (DCT) is a graph $\Gamma$ with $2^{N-1}$ nodes representing all possible developmental contexts. $\Gamma$ contains a directed edge $\alpha \to \beta \mid \alpha, \beta \in \{1, 2, .., 2^{N-1}\}$ whenever the developmental program encodes a transition from developmental context $\kappa_\alpha$ to $\kappa_\beta$. Since developmental contexts do not take number of cells of different cell types into account, we build $\Gamma$ using cell differentiation rules $Diff$ that encode differentiation and cell-death:

Let $\xi_\alpha$ be an N-length vector, such that

$$\begin{aligned}\xi_\alpha^i &= 1, \text{ for } \sum_{k=1}^{N}(\kappa_\alpha^k \times Diff_{\chi_\alpha}^{k,i}) > 0 \\ &= 0, \text{ otherwise}\end{aligned}$$

Then, there is an edge $\alpha \to \beta$ in graph $\Gamma$ whenever $\kappa_\beta^i = \xi_\alpha^i, \ \forall i \in \{1, 2, .., N\}$.

'Normal' development in the model begins with the progenitor, and follows a deterministic sequence of organ state transitions. In most cases, this set of transitions does not involve all $2^{N-1}$ possible developmental contexts. We call the DCT path from developmental context 0 (i.e., the progenitor) to the steady state the 'backbone' of the DCT.

### 7 Cell Differentiation Maps

Cell differentiation maps (CDM) represent the set of cellular differentiations that occurred during development and include information about the timing of first appearance of different cell types in the developing organ. Mathematically, a CDM is composed of a graph of cellular differentiations $G$ and an $N$-length vector $\tau$ that records the time of first appearance of the N-cell types.

The graph $G$ has $N$ nodes which represent different cell types. There is a directed edge from node $i$ to node $j$ if cell type-$i$ differentiated into cell type-$j$ at any time-step during development. We represent the graph $G$ using an $N \times N$ adjacency matrix $A$:

$$\begin{aligned}A^{i,j} &= 1, \text{ if} \sum_{t=0}^{t_{ss}} Diff_{\chi_t}^{i,j} > 0 \\ &= 0 \text{ otherwise.}\end{aligned}$$

And, the appearance time for any cell type-i is given by $\tau^i = t_i \mid (\Omega^i_{t<t_i} = 0)$ & $(\Omega^i_{t_i} > 0)$. To obtain the alignment of differentiation edges with the direction of developmental time, we define another $N \times N$ binary matrix $dir$:

$$\begin{aligned}dir^{i,j} &= 1 \text{ if } \tau^j - \tau^i \geq 0 \\ &= 0 \text{ if } \tau^j - \tau^i < 0\end{aligned}$$

We quantify the following properties of the CDM:

- Number of convergent differentiations: $\sum_{j=1}^{N}((\sum_{i=1}^{N} A^{i,j}) - 1)$.
- Number of *downward* convergent differentiations: $\sum_{j=1}^{N}\sum_{i=1}^{N}(A^{i,j} \times \delta_{dir^{i,j},1})$. Here, $\delta$ refers to the Kronecker delta function.

| N | Stable only | Expanding only | Shrinking only | Stable + Expanding | Stable + Shrinking | Expanding + Shrinking | All three dynamics |
|---|---|---|---|---|---|---|---|
| 3 | 0 | 0 | 0 | 11 | 0 | 0 | 2 |
| 4 | 22 | 89 | 0 | 200 | 0 | 1 | 9 |
| 5 | 868 | 3979 | 110 | 315 | 7 | 49 | 2 |
| 6 | 918 | 5382 | 69 | 29 | 0 | 10 | 0 |

Table. S 1: Developmental programs with distinct growth dynamics can map to the same CDM topology. Numbers indicate the number of CDM topologies produced by sets of programs with growth dynamics as indicated by column labels.

- Number of *backward* convergent differentiations: $\sum_{j=1}^{N}\sum_{i=1}^{N}(A^{i,j} \times \delta_{dir^{i,j},0})$. Note that all cyclic differentiations are necessarily *upward* edges, but *backward* edges can also be acyclic.

Many distinct developmental programs produce the same CDM topology, therefore, representing developmental programs as CDMs incurs information loss (Fig. S4A). In fact, even programs leading to disinct growth dynamics can map to the identical CDM, i.e., CDMs cannot differentiate between growth dynamics (Table 1). Nevertheless, there are statistical associations between growth dynamics and CDM topology; importantly, some CDMs are enriched for among the stable subset of sampled programs (Fig. S4B).

#### 7.1 Stable programs, whether fixed-points or limit cycles, are depleted for convergent differentiations

The CDMs that are differentially enriched among stable programs tend to have fewer convergent, especially backward convergent differentiations. In other words, they tend to be treelike and irreversible. This is true irrespective of whether the stable programs lead to fixed-points or limit cycle steady states (Fig. S5). When all programs are taken together, there is a sharp depletion of CDMs with 1 convergent differentiation; this is driven

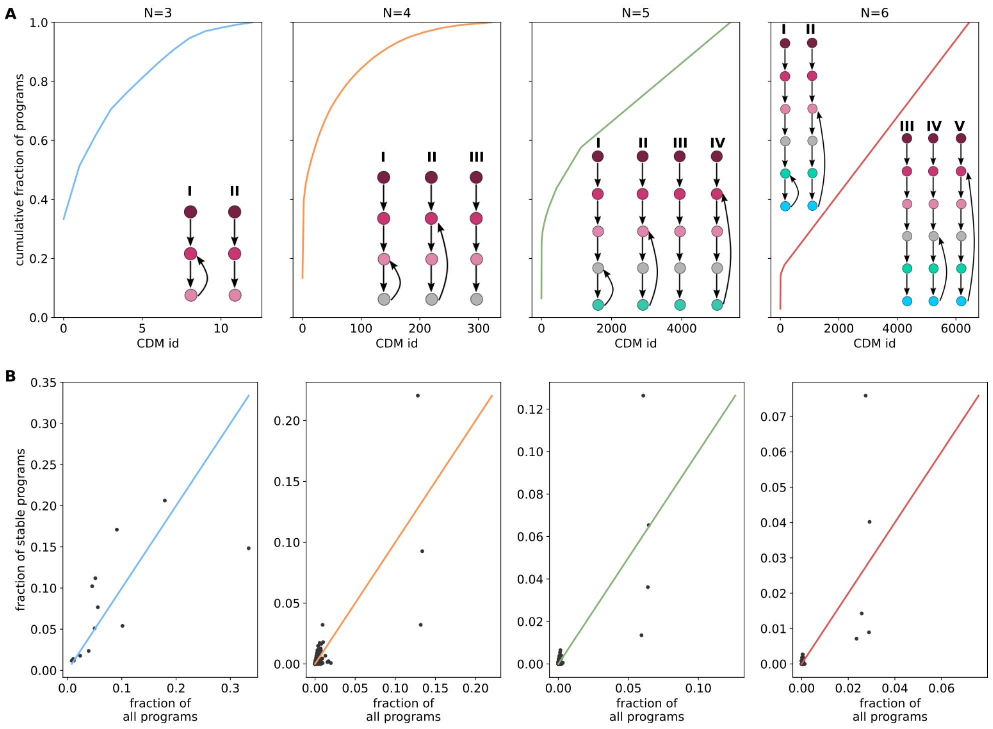


Fig. S4: Information loss in CDM representation. (A) Line plots show the frequency of distinct CDM topologies among sampled N=3,4,5,6 programs. Points on the x-axis represent CDM topologies ordered according to their popularity. The y-axis represents the cumulative fraction of sampled programs that map to each CDM topology. Insets: Most popular CDM topologies in order of popularity in sampled data. (B) CDM topologies enriched among stable programs. Each dot represents a distinct CDM topology. For each N, the x-axis indicates the fraction of non-trivial, valid programs that produce some CDM topology. The y-axis indicates the fraction of non-trivial, valid stable programs that produce this CDM topology. Dots above the colored lines are enriched among the stable subset of programs, and dots below this line are depleted among the stable subset of programs.

by a handful of exceptionally popular CDMs (Fig. S4A): a smaller fraction of stable programs as compared to all non-trivial programs produce these CDMs (i.e. they are depleted among stable programs), and these CDMs have an outsize effect when all CDMs are pooled together (Fig. S6, columns 1, 2). These exceptionally popular CDMs tend to be produced by expanding programs with limit cycle steady states; this is apparent in Fig. S5B, where there is a sharp depletion of CDMs with 1 convergent differentiation, while there is no such dip in Fig. S5A. There are no exceptionally popular CDMs among those that have higher number of convergent differentiations (Fig. S6, columns 3, 4).

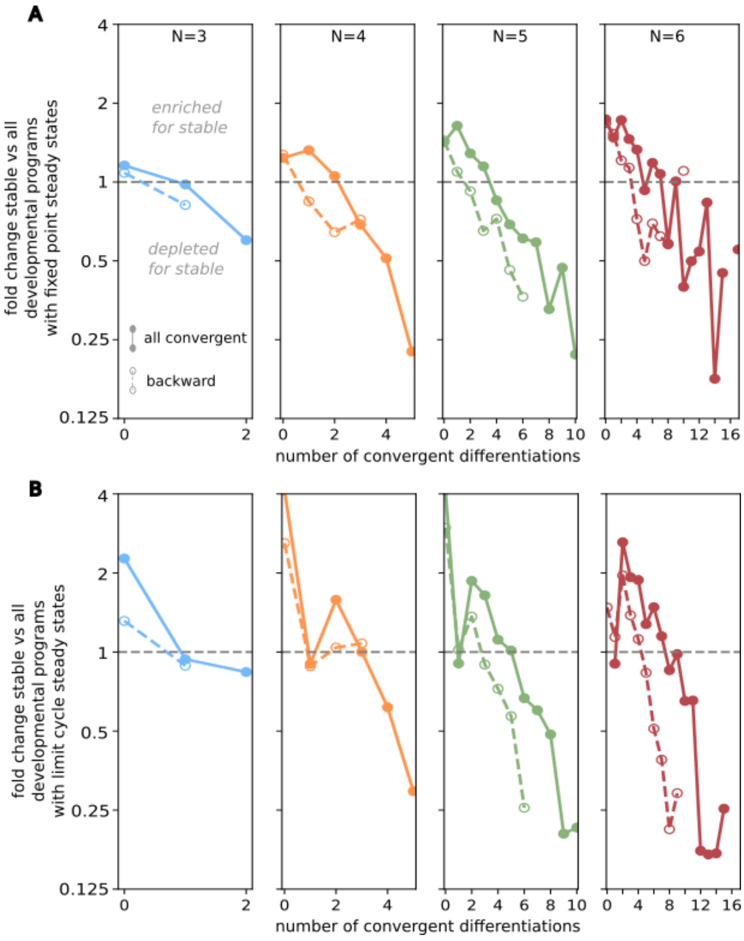


Fig. S5: Enrichment vs depletion of all convergent (solid) and backward convergent (dashed) differentiations in stable programs with $N = 3, \ldots, 6$ cell types with (A) fixed point steady states, (B) limit cycle steady states. Vertical axis shows the fold-change in the proportion of CDMs for stable vs all sampled developmental programs with a given number of convergent differentiations (horizontal arrow). Values $> 1$ indicate enrichment, values $< 1$ depletion. Stable programs, both fixed points and limit cycles, are enriched for zero convergent differentiations and strongly depleted for high number of convergent differentiations .

### 8 Mutations and perturbations

#### 8.1 Mutations to developmental programs

A mutation to the developmental program occurs in $Diff$ (differentiation and death rules). Let some mutation result in mutant rules $MutDiff$. $MutDiff$ is different from $Diff$ in the differentiation rule for a single cell type in a single developmental context. Let $\chi$ be a developmental context randomly chosen from the set $\{1, 2, .., 2^{N-1}\}$, and $i, j$ be a pair of cell-states randomly chosen from the set $\{2, 3, .., N\}$. $i$ and $j$ are allowed to be identical. Then,

$$\begin{aligned} MutDiff_{\chi}^{i,j} &= 0 \text{ if } Diff_{\chi}^{i,j} > 0 \\ &= \frac{1}{targets_{\chi}^{i} + 1}, \text{ if } Diff_{\chi}^{i,j} = 0 \end{aligned}$$

If the final steady state of a program remains unchanged despite a mutation, we call the program robust to this mutation. In this study, we simulate 100 independent mutations to each non-trivial, valid developmental program, and calculate the *mutation robustness index*: the fraction of mutations to which a program is robust. A majority of programs in our dataset were robust to mutations, irrespective of their growth dynamics (Fig. S7A, Fig. S9A). Mutational robustness was not strongly correlated with any of the three sampling parameters used in this study (Fig. S7B).

#### 8.2 Perturbations to steady state

We perturb the number of cells of a single cell type in a single phase of the steady state cycle.

Ablation/insertion of a cell type: Let $i$ be a cell type randomly chosen from the set $\{2, 3, .., N\}$, and $\chi$ be a developmental context randomly chosen from within the steady state cycle $\{\chi_{t_{ss}-p}, .., \chi_{t_{ss}}\}$. Let $Perturbed\Omega$ represent the perturbed organ. Then,

$$
\begin{aligned}
Perturbed\Omega_{\chi}^{i} &= 1 \text{ if } \Omega_{\chi}^{i} = 0 \\
&= 0, \text{ if } \Omega_{\chi}^{i} > 0
\end{aligned}
$$

If the final steady state of a program remains unchanged despite a perturbation, we call the program robust to this perturbation. In this study, we simulate 100 independent perturbations to each non-trivial, valid developmental program, and calculate the *perturbation robustness index*: the fraction of perturbations to which a program is robust. Programs in our dataset were fairly robust to perturbations, irrespective of their growth dynamics (Fig. S8A, Fig. S9B). Perturbational robustness is fairly strongly correlated with the sampling parameter for cell death $\phi$ (Fig. S8B).

### 9 Somatic mutations and cancer

A somatic mutation arises in a single cell in the steady state organ and leaves all other cells, including other cells of the same cell type untouched. In the model, we simulate a somatic mutation as the introduction of N new cell types, representing mutant cell types, to the developmental program. That is, for some cell type $i$, the

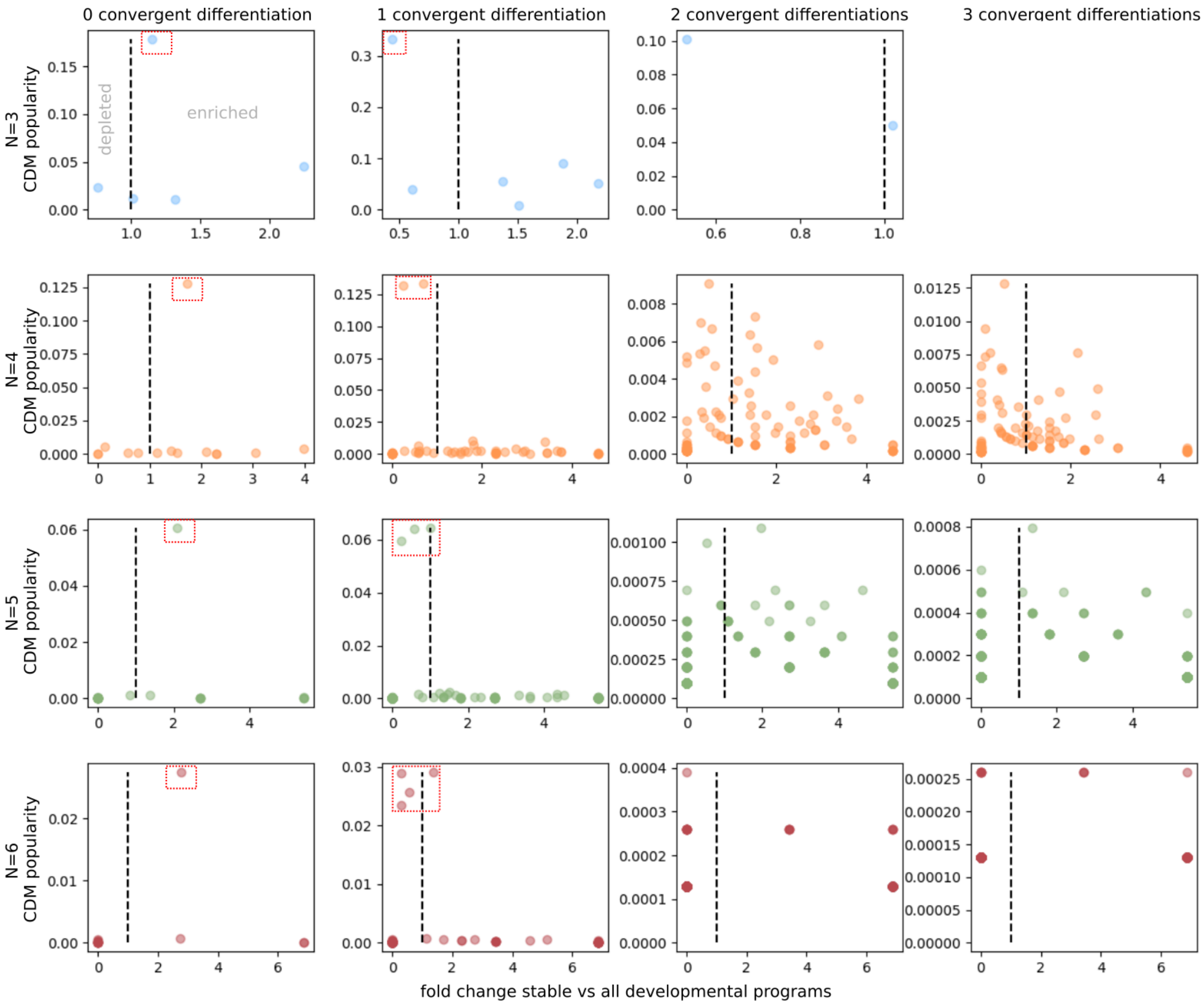


Fig. S6: Exceptionally popular CDMs drive depletion statistics. Each dot in these scatter plots represents a distinct CDM with a given number of convergent differentiations (columns) for programs with 3,4,5,6 cell types (rows). Horizontal axes represent the fold change in the fraction of stable versus all non-trivial programs that produce some CDM. Vertical axes represent CDM popularity; i.e., the fraction of sampled non-trivial programs that produce this CDM. Exceptionally popular CDMs are highlighted in red boxes.

corresponding mutant type is $i+N$. A randomly chosen cell in the steady state organ switches from its original cell type to its corresponding mutant type.

Somatic mutations cause a mutant cell to perceive itself to be in an incorrect context. Any given mutation is associated with a specific error in perception: A somatic mutation maps each context $k \in [0,1]^N$ to some randomly chosen (incorrect) context ${}^{sm}\kappa \in [0,1]^N$. The same somatic mutation can be incident on any cell of any cell type, however, the mutation does not damage the ability of the cell to perceive its own cell type: A mutant cell of type $i+N$ in some context $\kappa$ perceives itself to be in context ${}^{sm}_{i}\kappa$ where,

$$\begin{aligned} {}^{sm}_{i}\kappa^i &= 1 \\ {}^{sm}_{i}\kappa^j &= {}^{sm}\kappa^j \text{ for } j \neq i \end{aligned} \tag{1}$$

Let the somatic mutation occur in a cell of cell type $i$ when the organ is in a state $\Omega_m \in \{\Omega_{t_{ss}} - p,\ \Omega_{t_{ss}} - p+1, ..., \Omega_{t_{ss}}\}$, which is one of the phases of its steady state. This somatic mutation introduces a new mutant cell type be $i+N$ into the organ. That is, the organ state changes from being the N-length vector ${}_m\Omega$ to the 2N-length vector $\Omega'_m$, where

$$\begin{aligned} \Omega'^{i}_m &= \Omega^i_m - 1 \\ \Omega'^{i+N}_m &= 1 \\ \Omega'^{j}_m &= \Omega^j_m \quad \forall j \neq i \end{aligned}$$

Non-mutant cells do not perceive mutant cells any differently from their normal counterparts, and the developmental context for all normal cells $i|i \in [1,..,N]$ is given by:

$$\begin{aligned} \kappa^i_t &= 1 \quad if\ \Omega'^i_t + \Omega'^{(i+N)}_t > 0 \\ &= 0 \quad if\ \Omega^i_t + \Omega'^{(i+N)}_t = 0 \end{aligned}$$

While the context perceived by any mutant cells in the organ $i+N|i \in [1,..,N]$ is given by eq.(1).

Post somatic mutation, both mutant and non-mutant cells make cellular decisions (division, death, differentiation) according to the developmental contexts they perceive, as described in section ??: 'Updating the state of an organ'. The somatic mutation is inherited by all daughter cells in the cell lineage of the initial mutant.

The somatic mutation perturbs the organ from its original steady state and leads to a new steady state. We call this somatic mutation cancerous if the steady state of the somatic mutant is of the Expanding type. If the

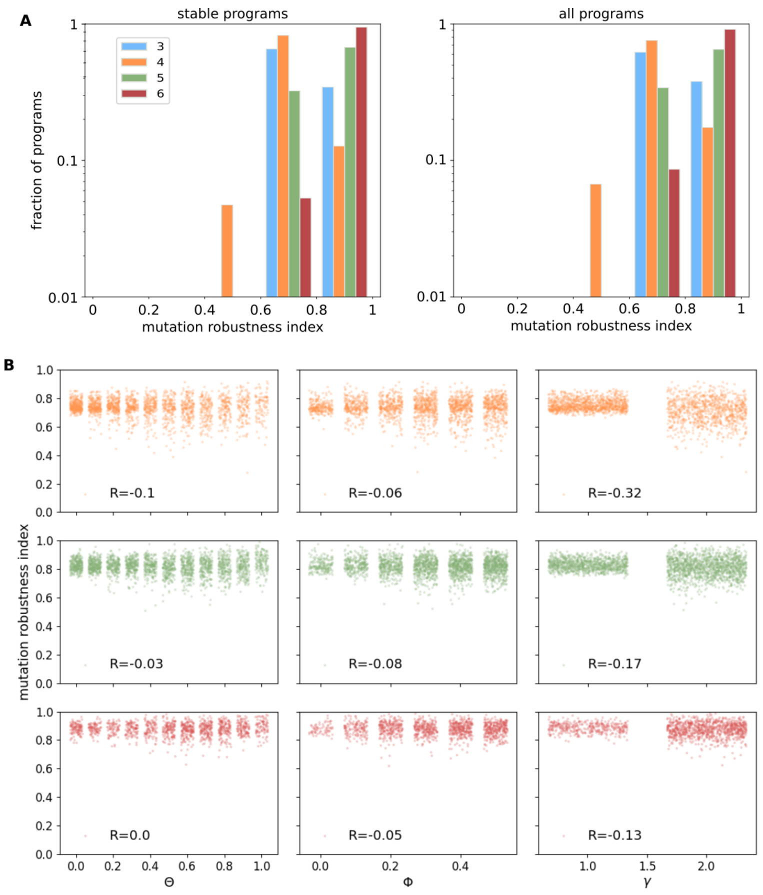


Fig. S7: Mutational robustness among sampled programs. (A) Histograms of mutational robustness index for (left) stable programs, (right) all non-trivial, valid programs. (B) Scatter plots showing correlations between model sampling parameters and mutation robustness index for (top row) N=4, (middle row) N=5, (bottom row) N=6.

final steady state of a program remains stable despite a somatic mutation, we call the program robust to this somatic mutation. In this study, we simulate 100 independent somatic mutations to each stable developmental program, and calculate the *cancer robustness index*: the fraction of somatic mutations to which a program is robust. Programs in our dataset were fairly robust to cancer(Fig. S10A). Cancer robustness is strongly negatively correlated with the sampling parameter for cell division $\theta$, positively correlated with the cell death parameter $\phi$ and negatively correlated with the heterogeneous differentiation parameter $\gamma$ (Fig. S10B).

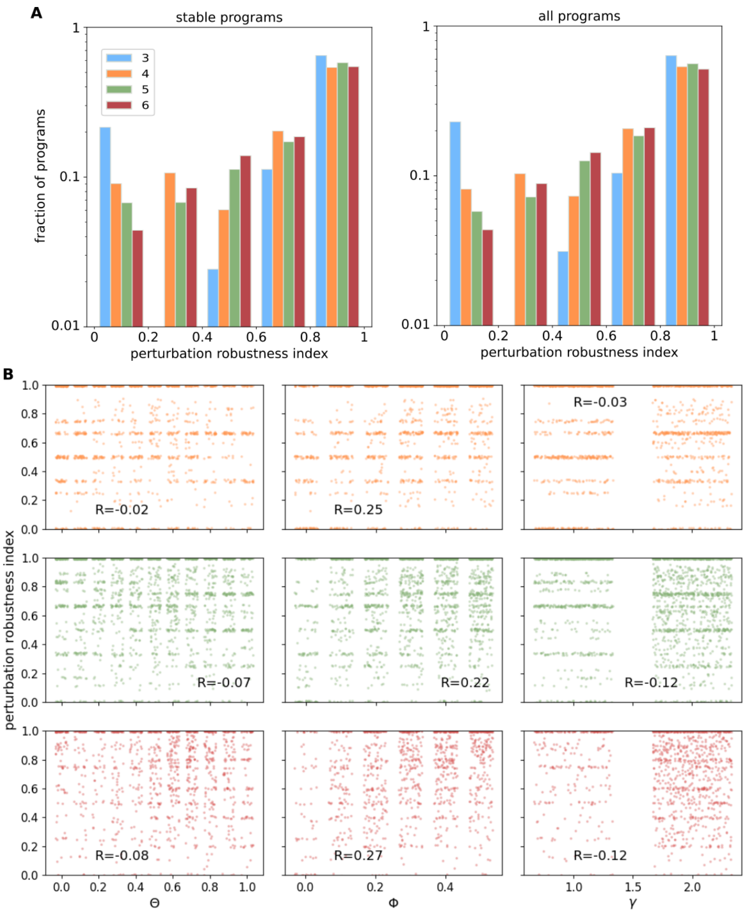


Fig. S8: Perturbational robustness among sampled programs. (A) Histogram of perturbation robustness index for (left) stable programs, (right) all non-trivial, valid programs. (B) Scatter plots showing correlations between model sampling parameters and perturbation robustness index for (top row) N=4, (middle row) N=5, (bottom row) N=6.

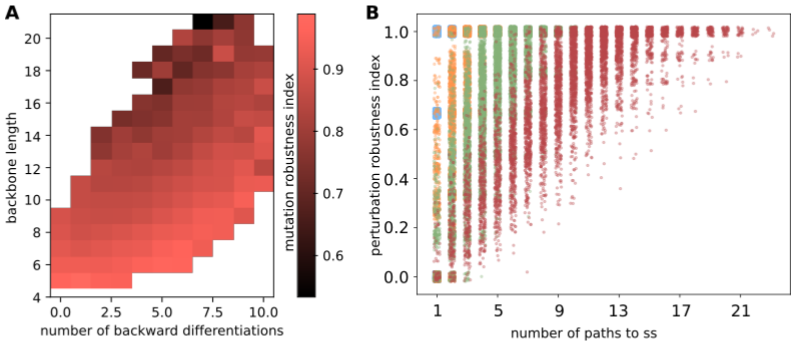


Fig. S9: Mutational and perturbational robustness across all non-trivial, valid developmental programs. (A) Heatmap representing the dependence of mutational robustness of N=6 developmental programs on DCT and the CDM topologies. Rows represent developmental trajectory length. Columns represent the number of *backward* differentiations in the CDM. Robust developmental programs have smaller trajectory lengths and fewer *backward* differentiations. (B) Each dot in the scatter plot represents a distinct developmental program. The x-axis represents number of DCT paths to the *backbone*. The y-axis represents the perturbation robustness index. Developmental programs with high perturbation robustness have more paths to the backbone; i.e., they are canalized.

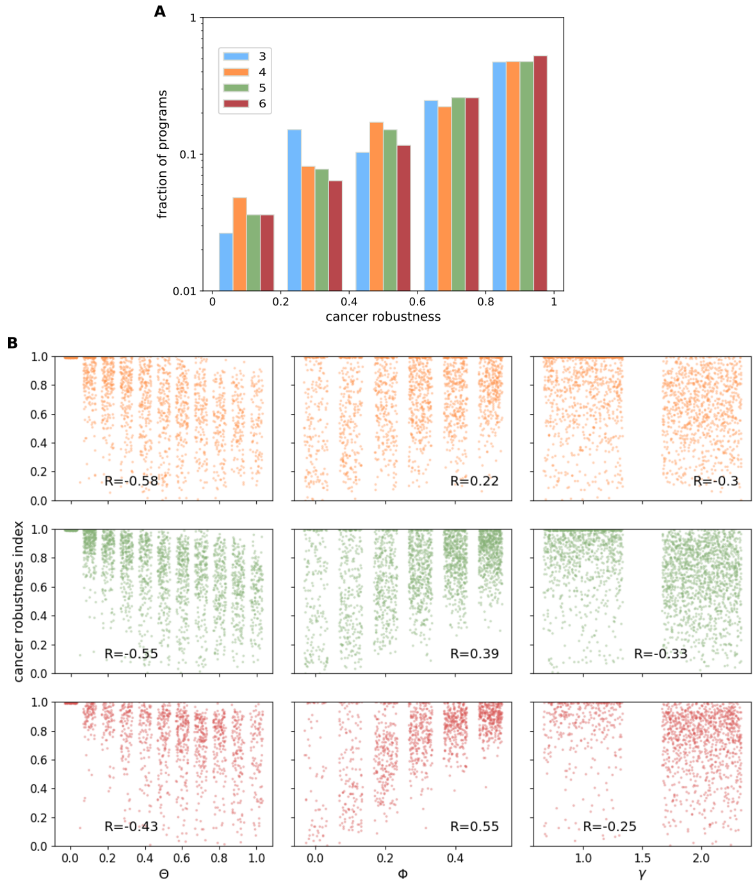


Fig. S10: Cancer robustness among stable programs. (A) Histogram of cancer robustness index for stable programs. (B) Scatter plots showing correlations between model sampling parameters and cancer robustness index for (top row) N=4, (middle row) N=5, (bottom row) N=6.
